# The protein phosphorylation landscape in photosystem I of the desert algae *Chlorella sp*

**DOI:** 10.1101/2023.10.01.560385

**Authors:** Guy Levin, Michael Yasmin, Tomasz Pieńko, Nitsan Yehishalom, Fabian Glaser, Gadi Schuster

## Abstract

- The phosphorylation of photosystem II (PSII) and its antenna (LHCII) proteins has been extensively studied and its involvement in state transitions and PSII repair is well known. Yet, very little is known about the extent and functions of phosphorylation of photosystem I (PSI) and its antenna (LHCI) proteins.
- Here, two proteomics methods were applied to generate a detailed map of the phosphorylation sites of the PSI-LHCI proteins in *Chlorella ohadii* cells that were grown under low- or extreme high-light intensities (LL and HL). Furthermore, we analyzed the content of oxidized tryptophans in these cell types to estimate light-induced oxidative damage to PSI-LHCI.
- Our work revealed the phosphorylation of 11 out of 22 PSI-LHCI subunits. The analyses detected extensive phosphorylation of the LHCI subunits lhca6 and lhca7. Other PSI-LHCI subunits were phosphorylated to a lesser extent. Additionally, we show the accumulation of oxidatively damaged tryptophans in the psaD subunit of PSI of HL-grown *C. ohadii*.
- The significant phosphorylation of lhca6 and lhca7 suggests a physiological role during photosynthesis, possibly by altering light-harvesting characteristics and binding of other LHCI subunits. Moreover, we show that psaD is susceptible to photodamage while LHCI is protected from ROS under HL.

## Introduction

Thylakoid protein phosphorylation was discovered decades ago (Bennett, 1977). Ever since, its importance in the regulation of photosynthesis under different light conditions has been demonstrated in many contexts (Allen, 1992; Jonwal *et al*., 2022). The most studied process is the phosphorylation of photosystem II (PSII) and light-harvesting complex II (LHCII) subunits. The core PSII proteins are phosphorylated in response to light stress, which initiates the disassembly of the damaged PSII (Aro *et al*., 1992; Tikkanen *et al*., 2008). This is a crucial step in the photosynthetic PSII repair cycle and the prevention of photoinhibition, a high-light (HL)-induced process that leads to the inhibition of photosynthesis (Schuster *et al*., 1988; Adir *et al*., 2005; Bassi & Dall’Osto, 2021). Phosphorylation of LHCII subunits mediates state transitions, during which some subunits are transferred from the PSII to the PSI complex, which subsequently enhances its absorbance cross-section and ensures balanced energy uptake between the two photosystems (Chow *et al*., 1981; Goldschmidt-Clermont & Bassi, 2015; Nawrocki *et al*., 2016). In contrast, the phosphorylation of PSI-LHCI subunits has been far less studied and its physiological role remains unknown (Grieco *et al*., 2016; Longoni & Goldschmidt-Clermont, 2021; Younas *et al*., 2023). Previously detected phosphor-sites were found in the PSI subunits of psaC, psaD, and psaE (Reiland *et al*., 2009; Wang *et al*., 2014; Spät *et al*., 2015), which are located in the stromal side of PSI where they form the binding site of ferredoxin (Fd) (Caspy *et al*., 2020). There, these subunits mediate electron transfer between PSI and Fd NADPH reductase (FNR), and thereby, between the photosynthetic light reactions and CO_2_ assimilation in the Calvin-Benson-Bassham cycle. Additionally, phosphor-sites were identified in psaH, psaG, psaK, and psaL (Reiland *et al*., 2011; Roitinger *et al*., 2015; Younas *et al*., 2023), which are involved in the binding of LHCII during state transitions. It is suggested that psaG and psaH phosphorylation stabilizes the PSI-LHCI-LHCII complexes during state II (Younas *et al*., 2023).

Recently, we found that when acclimated to low light (LL), *Chlorella ohadii*, an extremely light-tolerant desert green alga (Treves *et al*., 2016; Levin *et al*., 2021; Levin & Schuster, 2023), has only limited state transitions, while when acclimated to HL, this process is absent (Levin *et al*., 2023). Accordingly, PSII-LHCII protein phosphorylation in thylakoids of state II-induced *C. ohadii* cells is negligible compared to other algae and plants. Phosphorylation of the PSI subunit psaE was detected in both LL *C. reinhardtii* and *C. ohadii* cells, but not in HL *C. ohadii*, suggesting possible light regulation of PSI-LHCI phosphorylation (Levin *et al*., 2023). This discovery prompted an investigation of the extent, location, and possible roles of PSI-LHCI protein phosphorylation in LL- and HL-grown *C. ohadii* cells.

This work presents a detailed map of protein phosphorylation of PSI-LHCI in LL- and HL-grown *C. ohadii*, including novel phosphor-proteins that were not previously described. LHCI subunits lhca6 and lhca7 are substantially and differently phosphorylated under different light conditions, suggesting the involvement of phosphorylation in light harvesting and binding of other LHCI subunits. Additionally, we show that many tryptophans of PSI-LHCI subunits are oxidized, a sign of photodamage, but only psaD displays a significant increase in tryptophan oxidation in the PSI complex of HL-grown cells. Surprisingly, LHCI subunits accumulate less oxidative damage in cells grown in HL conditions, suggesting a protective mechanism that is induced during their adaptation to HL.

## Materials and Methods

### PSI-LHCI isolation

Log phase *C. ohadii* cells were grown in growth chambers (28 °C) in tris-acetate-phosphate (TAP) medium under LL (50 µmol photons m^-2^ s^-1^) or HL (2000 µmol photons m^-2^ s^-1^) conditions for at least 6 generations. Cells were maintained below O.D_750nm_ = 0.7 to prevent self-shading. Cells were harvested, washed twice with sucrose-Tris-tricine-NaCl_2_ (STN), and then broken under 50 PSI with a MicroFluidizer. Samples were centrifuged (12000 rpm, 10 min, 4 °C), the supernatant was collected and thylakoid membranes were pelleted via ultra-centrifugation (50000 rpm, 2 h, 4 °C) and then resuspended in 25mM MES (pH 6.5) to a chlorophyll concentration of 0.2 (HL) or 0.4 (LL) mg/ml. Thylakoid membranes were solubilized 15 min with 1% n-dodecyl-α-D-maltopyranoside (α-DDM) on ice, in the dark and insolubilized material was pelleted (10000 rpm, 10 min, 4 °C). The supernatant was collected and loaded on a sucrose density gradient (20 mM MES pH 6.5, 0.02% α-DDM). Individual photosynthetic complexes were fractionated by ultra-centrifugation (25,000 rpm, 20 h, 4 °C). The PSI fraction from three biological repeats of LL and HL cells was collected and analyzed via mass spectrometry.

### Immunoblot of PSI-LHCI phosphorylated peptides

Isolated PSI-LHCI complexes were fractionated by denaturing SDS-PAGE and blotted to nitrocellulose membrane. The membranes were blocked and incubated overnight with antibodies against phosphorylated Serine and Threonine. Membranes were then washed and incubated with secondary antibody for 1 hour. Chemiluminescence signals were detected using Vilber Fusion Pulse (Collégien, France).

### Mass spectrometry sample preparation and proteolysis

The PSI-LHCI protein samples were brought to 8.5M Urea, 100 mM Ammonium bicarbonate, and 10 mM DTT (Omni TH Tissue Homogenizer). About 200 μg proteins from each sample were reduced with 10 mM DTT (60°C for 30 min), modified with 40 mM iodoacetamide in 400 mM ammonium bicarbonate (in the dark, room temperature for 30 min), and digested overnight at 37° C in 1.5M Urea with modified trypsin (Promega) at a 1:50 enzyme-to-substrate ratio. An additional second trypsinization was done for 4 hours, the samples were then acidified with 0.1% TFA and desalted with C18 Oasis columns. 20 μg of peptides was taken for proteomics analysis. The rest were re-suspended in 40% Acetonitrile (ACN), 6% TFA (Trifluoroacetic acid) and enriched for phosphopeptides using titanium dioxide (TiO 2) beads.

### Phosphopeptides enrichment using titanium dioxide (TiO2) beads

The Titanium beads were pre-washed (80% ACN, 6% TFA) and mixed with the peptides for 10 minutes at 37 °C, washed with 30% ACN with 3% TFA and 80% ACN, with 0.1%TFA. Bound peptides were eluted with 20% ACN and 325 mM Ammonium Hydroxide followed by 80% ACN containing 325 mM Ammonium Hydroxide. The resulting peptides were desalted using C18 tips and analyzed by LC-MS-MS.

### Mass spectrometry analysis

Untreated and phospho-enriched peptides were resolved by reverse-phase chromatography on 0.075 × 180-mm fused silica capillaries (J & W) packed with Reprosil reversed phase material (Dr Maisch GmbH, Germany). The peptides were eluted with different concentrations of Acetonitrile with 0.1% of formic acid: a linear 60 minutes gradient of 6 to 34% acetonitrile followed by a 15-minute gradient of 34 to 95% and 15 minutes at 95% acetonitrile with 0.1% formic acid in water at a flow rate of 0.15 μl/min. Mass spectrometry was performed by Exploris 480 mass spectrometer (Thermo) in a positive mode (m/z 300–1500, resolution 60,000 for MS1 and 15,000 for MS2) using repetitively full MS scan followed by collision-induced dissociation (HCD, at 27 normalized collision energy) of the 20 most dominant ions (>1 charges) selected from the first MS scan. A dynamic exclusion list was enabled with an exclusion duration of 20 seconds.

### Data analysis

Raw data were processed using MSFragger version 3.7 (Kong *et al*., 2017) via FragPipe version 19.1 (https://fragpipe.nesvilab.org/). As the proteome of *C. ohadii* was not yet available during this work, all of Chlorella protein sequences (Tax ID=3701) downloaded from UniProt, were used for protein identification (20,973 sequences; July, 2019). These sequences were supplemented with sequences of known contaminants and decoy sequences via FragPipe. All searches included (unless specified otherwise) enzymatic cleavage with “stricttrypsin” and 2 missed cleavages and clip of N-term methionine. Cysteine Carbamidomethyl (+57.02146) was set as fixed modification and methionine oxidation (+15.994915), and protein N-terminal acetylation were set as variables modifications. Phosphorylation identification of after phosphopeptide enriched samples or unenriched samples were done using Fragpipe “LFQ-Phospho” workflow using its default settings that include phosphorylation of serine, threonine, or tyrosine as variable modification and PTM site localization with PTMProphet and label-free quantification using IonQuant with match-between-runs. Identification of tryptophan oxidation and semi-tryptic peptides was performed using FragPipe’s “LFQ-MBR” workflow. The workflow default settings were used with the additions of tryptophan oxidation (+15.994915) and deoxidation (+31.989829) as variable modifications for identifying tryptophan oxidation. For identifying semi-tryptic cleavages, we set the cleavage type to “semi”.

### Parametrization of the iron-sulfur clusters for molecular dynamics analysis

The iron-sulfur clusters coordinated by cysteines were modeled bonded using the MCPB.py software (Li & Merz, 2016). To obtain their bonded parameters, we used a “sidechain model” that included the iron-sulfur part and the sidechains of the cysteines, each terminated by a methyl group. The sidechain models of [4Fe-4S] and [2Fe-2S] clusters were optimized at unrestricted B3LYP* (B3LYP with 15% Hartree-Fock exchange) with the def2-TZVP basis set. B3LYP* has been shown to provide an accurate description of the electronic structure of iron-sulfur clusters (Salomon *et al*., 2002; Benediktsson & Bjornsson, 2022). An initial guess was generated based on a fragmentation approach, followed by a wavefunction’s stability calculation. For [4Fe-4S], eight fragments were defined, four of which contained an iron ion and methyl sulfide, and the other four were only sulfides. The former fragments had the following charge/multiplicity: 2/-6, 2/6, 1/-5, 1/5, while the latter had a charge of −2 and multiplicity of 1, giving a total molecular charge of −2 and multiplicity of 1. For [2Fe-2S], four fragments were created, among which two included an iron ion and two methyl sulfides and the other two were sulfides. The former fragments had a charge/multiplicity of 1/-6 and 1/6, while the latter had a charge of −2 and multiplicity of 1, yielding a total molecular charge of −2 and multiplicity of 1. Calculations of the spin densities of the fragments confirmed that a correct wavefunction representing an antiferromagnetic spin-coupled state (Noodleman *et al*., 1995) was obtained. At the optimized geometry, vibrational frequencies analysis was performed to obtain bond and angle force constants using the Seminario method (Seminario, 1996). As in the original approach, the force constants of the dihedral angles were neglected. Van der Waals parameters for the iron ions optimized for OPC water in the IOD set of 12-6 model were used (Li *et al*., 2021), and for sulfides they were assigned using GAFF2. The atomic charges of the iron-sulfur clusters and ligating cysteines were computed using a “large model” that consisted of the iron-sulfur component and cysteines with their backbones capped with acetyl (ACE) and N-methyl (NME) at the N and C termini, respectively. To assure the antiparallel, spin-coupled state of the wavefunction in the large model, fragmentation of the molecule and wavefunction’s stability calculations were carried out analogously to the sidechain model. The large models of [4Fe-4S] and [2Fe-2S] were optimized at unrestricted B3LYP* with the def2-SVP basis set. At the optimized structure, a single-point energy calculation was performed using the def2-TZVP basis set to generate the electrostatic potential. This was subsequently used by antechamber (AmberTools23) (www.ambermd.org) to derive the atomic RESP charges (Bayly *et al*., 1993) of the iron-sulfur clusters and coordinating cysteines. As recommended by (Peters *et al*., 2010), the charges of the backbone heavy atoms of cysteines were restrained to the values in the ff19SB protein Amber force field (Tian *et al*., 2020). All the quantum chemical calculations were performed using Gaussian 09 (www.gaussian.com).

### Molecular dynamics system build

We used the structure of Photosystem I from *Chlorella ohadii* (PDB code: 6ZZX) (Caspy *et al*., 2021) to model the PsaC, PsaD and PsaE proteins and their corresponding [4Fe-4S] clusters. Then, we employed AlphaFold-multimer-v3 (Evans *et al*., 2021) to create a hetero tetramer of the same proteins complexed with ferredoxin which is missing in the 6ZZX PDB structure. Finally, we aligned the AlphaFold highest ranked model of ferredoxin to the structure of ferredoxin from *Pisum sativum* (PDB code: 6YAC) (Caspy *et al*., 2020) and removed from the AlphaFold hetero-tetramer PsaC, PsaD and PsaE proteins remaining only the ferredoxin model, and added the [2Fe-2S] cluster from the 6YAC PDB structure to complete the system. Two systems, with unphosphorylated and phosphorylated Ser63, were built with tleap (AmberTools23). The proteins’ complex was solvated with a 12 Å layer of OPC (Izadi *et al*., 2014) water molecules in a truncated octahedron box. The system was neutralized and supplied with 0.15 M NaCl (Sengupta *et al*., 2021). For parametrization of the proteins, the ff19SB Amber force field was used (Tian *et al*., 2020).

### Molecular dynamics (MD) simulations protocol and analysis

The equilibration protocol consisted of an initial minimization and several steps of heating, and a gradual reduction of initial positional restraints. The equilibration consisted of a total of 8 ns of MD with a time step of 1 fs (stages 1–8) and 2 fs (stage 9). First, water and hydrogen atoms were energy-minimized (stage 1), followed by 1 ns of heating, using restraints in NVT ensemble (stage 2). Then, 1 ns of MD in NPT ensemble was carried out to adjust the density of water with the fully restrained solute (stage 3). Subsequently, 1 ns of MD with lower restraints in NPT (stage 4) was conducted. Then, a second energy minimization of the side chains was performed (stage 5), followed by three stages of 1 ns MD in NPT with decreasing restraints on the backbone (stages 6–8). Finally, a 2 ns unrestrained run in NTP (stage 9) was performed. The MD run was then continued in NPT for 200 ns to reach the stabilization of the solute’s RMSD. For each of the unphosphorylated and phosphorylated states, we performed 4 independent MD production runs of 250 ns each. Hydrogen-mass repartitioning was used to increase the time step to 4 fs. Periodic boundary conditions and Ewald sums (grid spacing of 1 Å) were used to treat long-range electrostatic interactions. All the MD simulations were run using pmemd.cuda (Amber 22). During the production runs, trajectory files were saved every 10 ps. All the analyses, including measurements of distances, RMSD, and interactions, were performed using cpptraj (AmberTools23) (www.ambermd.org). For short-range donor-acceptor interactions, we applied criteria of the donor–acceptor distance cutoff of 3.2 Å and a 150° cutoff for the donor-hydrogen-acceptor angle. The figures visualizing structures from MD were prepared using VMD (Humphrey *et al*., 1996). The plots were generated using gnuplot 5.2.

## Results

### lhca6 and lhca7 are highly phosphorylated at amino acids located in their interface with other LHCI subunits

To analyze the extent and differences of phosphorylation of PSI-LHCI proteins in LL- and HL-grown *C. ohadii* cells, thylakoids were solubilized with a detergent and the photosynthetic complexes were fractionated by centrifugation in a sucrose density gradient (SDG) (Fig. S1). The PSI-LHCI fractions were collected, digested with trypsin, and then were split into two samples. One sample underwent phosphopeptide enrichment prior to mass spectrometry (MS) analysis. The other was directly subjected to MS without phosphopeptide enrichment (Fig. 1). By combining the results of these two analyses, a comprehensive landscape of phosphorylated PSI-LHCI proteins was obtained, as each approach missed certain phosphorylations. In both samples, phosphorylated threonine, serine, and tyrosine were detected. Most notably, when compared to other PSI-LHCI proteins, the lhca6 and lhca7 subunits accounted for the majority of identified phosphorylated peptides. Across six biological repeats (comprising 3 LL and 3 HL samples), 78 phosphorylated peptides were detected for lhca6 and 47 for lhca7. For context, psaC had 20 identified phosphorylated peptides, while only a handful were detected in other subunits (Table 1). It is worth noting that this doesn’t automatically suggest that lhca6/7 are the most phosphorylated among PSI-LHCI proteins. Detecting phosphorylation in other subunits might be more challenging with MS because of their specific characteristics. Still, the data strongly indicated a significant phosphorylation of amino acids in these subunits. Interestingly, lhca6 showed more extensive phosphorylation in PSI-LHCI of HL-grown cells, whereas lhca7 predominantly exhibited phosphorylation in PSI-LHCI of LL-grown cells. This implies that PSI-LHCI phosphorylation is modulated by light intensity during cell growth. The phosphorylated amino acids in lhca6 and lhca7 were positioned at their interface with the LHCI subunits lhca4 and lhca8, respectively. This positioning suggests a potential role in adjusting the stability and light-harvesting capability of the PSI-LHCI complex (Fig. 2a). Immunoblot analysis of isolated PSI-LHCI complexes using anti-Phospho-Ser/Thr antibody yielded a notably stronger signal in HL compared to LL PSI-LHCI. This signal was mainly from a band located at approximately 25 kDa, consistent with the size of LHCI subunits (Fig. 2b). Because most of the phosphorylated peptides belonged to lhca6 and were more abundant in HL PSI-LHCI, it can be concluded that the signal represents lhca6 alone, or perhaps lhca6 and lhca7 together. In addition, while MS successfully detected phosphorylation of other PSI subunits, immunoblotting analysis detected negligible amounts of additional phosphorylated subunits, suggesting that while lhca6 and lhca7 are significantly phosphorylated, phosphorylation levels of other subunits are relatively low and only exist in a small population of the proteins, perhaps during damage, assembly, or disassembly of PSI.

**Figure 1.**
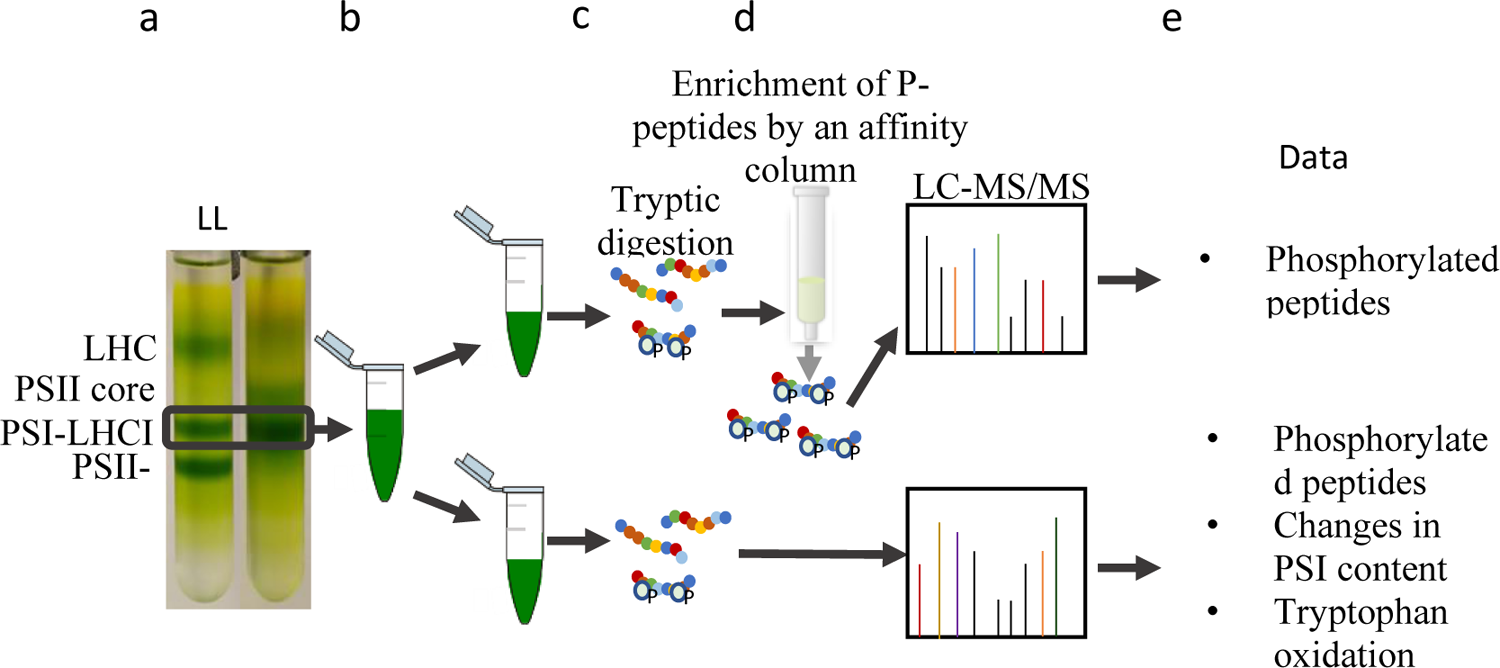
The workflow of the experiments in which the landscape of PSI protein phosphorylation has been analysed. *C. ohadii* cultures were grown for 24 h in low light (LL) or high light (HL) illumination. (a) Thylakoids of LL and HL cells were solubilized and photosynthetic complexes were fractionated using sucrose density gradient. (b) LL and HL PSI-LHCI complexes were collected and the proteins were cleaved by trypsin to generate peptides (c). (d) Half of the peptides underwent enrichment for phosphorylated peptides prior to LC-MS/MS analysis by using a phospho-peptide binding column (d top). The other half was loaded directly to the LC-MS/MS (d bottom). (e) Unprocessed data of each procedure was analysed to detect phosphorylation of Serine, Threonine, and Tyrosine in PSI-LHCI subunits. In addition, changes in the peptide contents of PSI-LHCI and oxidation of tryptophan residues were analysed in the peptides sample that was not enriched for phosphoproteins by the column.

**Figure 2.**
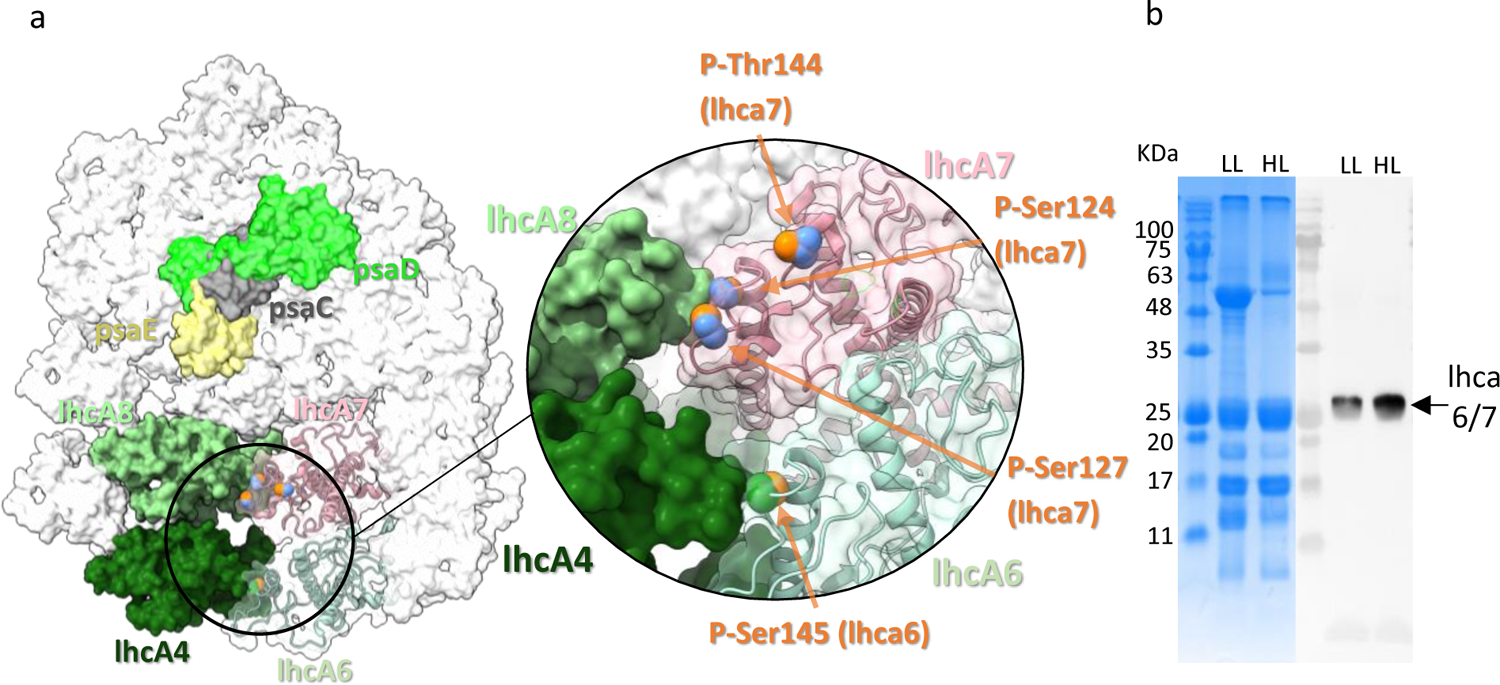
lhcA6 and lhcA7 are heavily phosphorylated at their interface with lhcA4 and lhcA8, respectively. (a) PSI-LHCI complex of *C. ohadii* (adapted from PDB: 6ZZX). Detected phosphorylation sites in lhca6 and lhca7 are indicated by orange orbs. Blue base indicates accumulation in LL and green base indicates accumulation in HL. psaC, psaD, and psaE are shown for context. (b) Immunoblot analysis using antibodies against p-Ser/Thr of isolated PSI-LHCI complex from HL and LL-grown *C. ohadii*. The Coomassie blue-stained SDS-PAGE is shown on the left panel and the immunoblot is on the right panel.

**Table 1.**
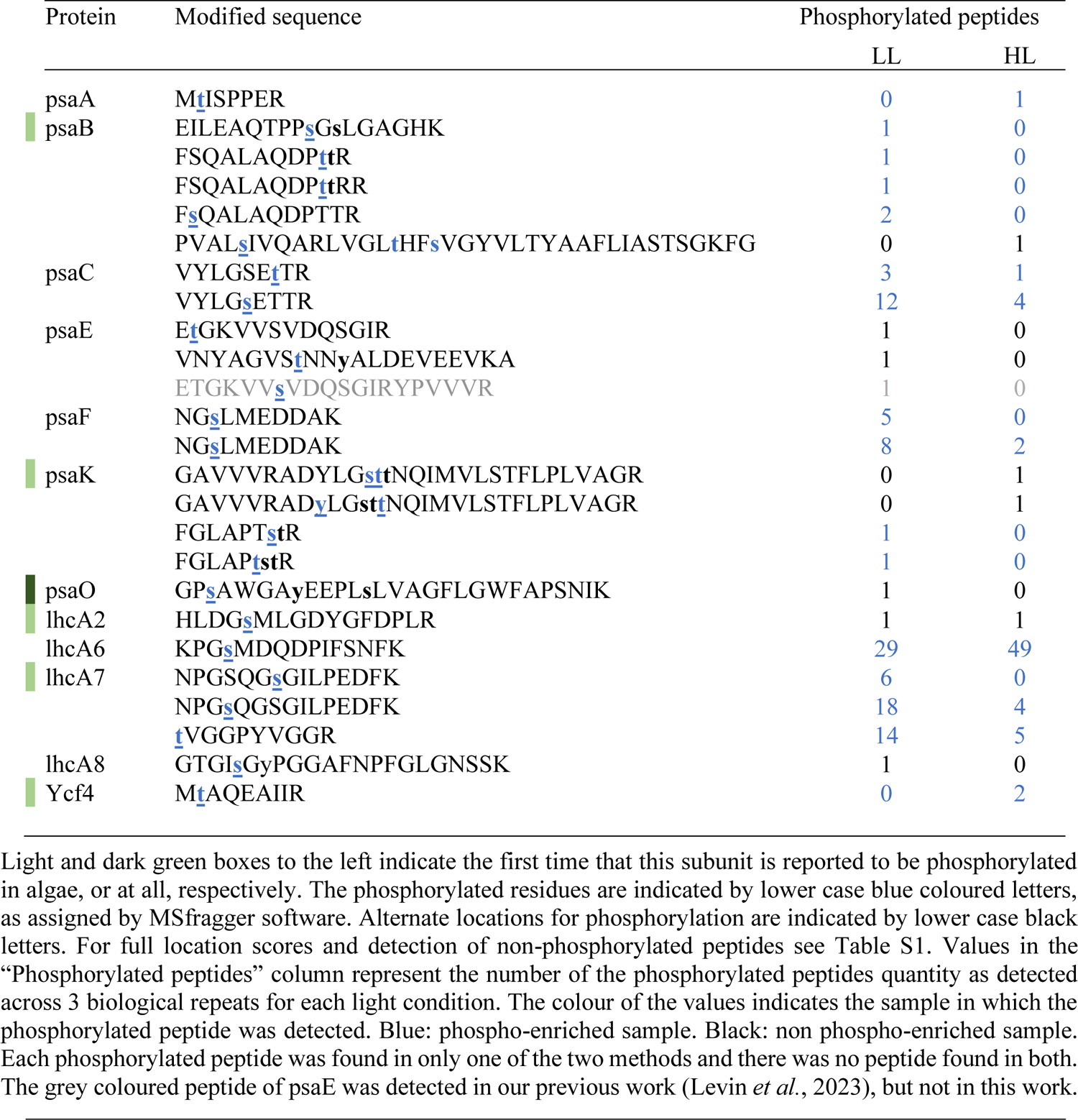
PSI-LHCI proteins are differentially phosphorylated in LL and HL-grown *C. ohadii* cells.

### Many PSI-LHCI phosphoproteins are phosphorylated at low levels, and some accumulate differentially in HL and LL-grown cells

Analyzing both total (non-enriched) and phosphopeptide enriched samples proved to be beneficial as several phosphorylated peptides that were detected in one sample, were not detected in the other (Table 1). Analysis of isolated PSI-LHCI complexes rather than thylakoid membranes or whole-cell extracts provided a more detailed phospho-sites map of PSI-LHCI in LL- and HL-grown *C. ohadii* and identified phosphoproteins that were not previously detected in green algae or plants (Fig. 3 and Tables 1, Table S1 and Table S2). Out of the 14 proteins of the PSI core complex that were detected in our MS analysis (Fig. 4 and Table S3), 7 were found to be phosphorylated (Table 1 and Fig. 3), including psaO, a phosphorylation which was not reported before in plants or algae. Phosphorylated psaO was only found in non-phosphor-enriched samples, which may explain why it was missed in previous analyses. Of the 8 detected antenna LHCI subunits, four were found to be phosphorylated. In addition to psaO, phosphorylation of psaB, psaK, lhcA2, and lhcA7 were identified here for the first time in green algae. Furthermore, phosphorylation of amino acids in psaA, psaC, psaE, psaF, lhcA6, and lhcA8 was demonstrated, as previously described by others. While it was reported in other organisms (Roitinger *et al*., 2015; Younas *et al*., 2023), no phosphorylation of psaH, psaL, and psaG was detected in *C. ohadii*. These subunits function in the binding of LHCII or stabilizing the complex during state transitions, a process that is considerably reduced in *C. ohadii,* which may explain the absence of these phosphorylations (Levin *et al*., 2023). Additionally, phosphorylation of ycf4, a crucial enzyme for PSI assembly, was detected. Of special interest are the identified phosphorylations of psaE and psaC (Fig. 2 and Table 1), which, together with psaD, form the binding site of Fd to PSI (Fig. 5a) (Caspy *et al*., 2020).

**Figure 3.**
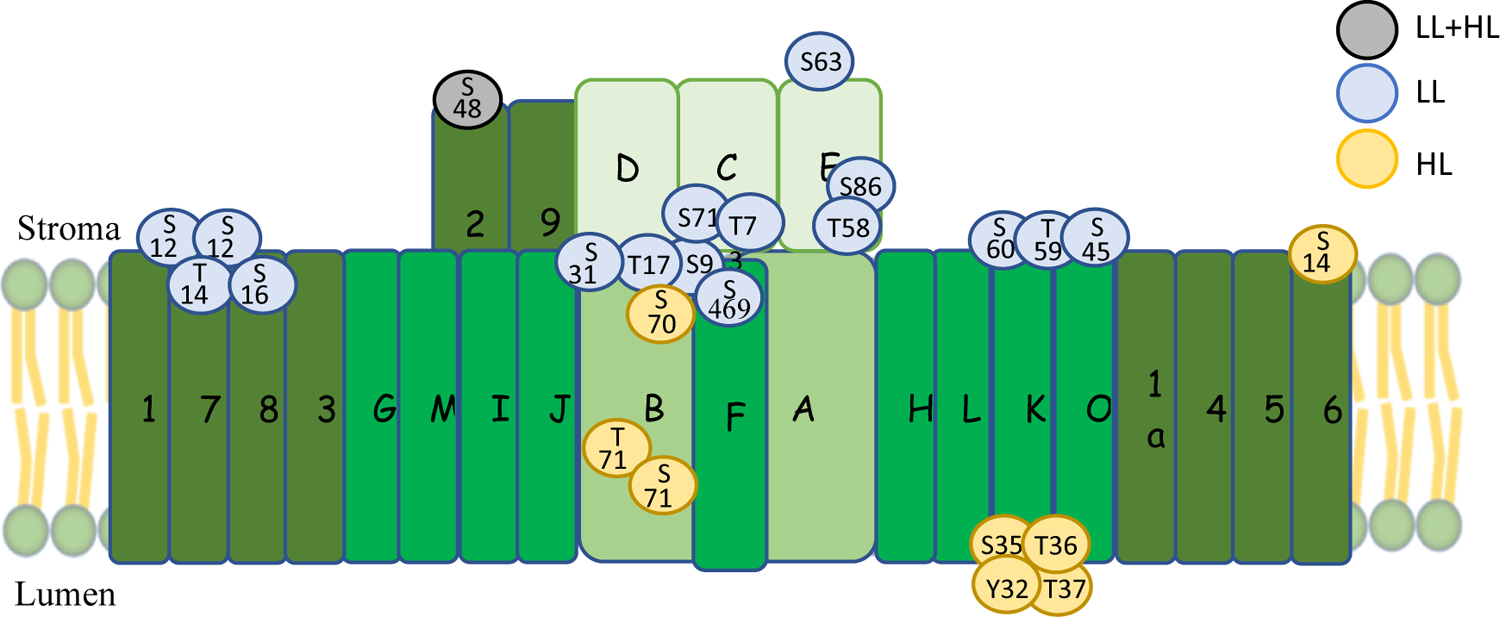
The landscape of PSI protein phosphorylation in LL and HL-grown cells. Blue and yellow balls represent phosphorylated amino acids that were detected in LL and HL-grown cells, respectively. A grey ball represents the amino acid that maintains a similar phosphorylation level in both LL and HL-grown cells. Letters and numbers within the balls represent the phosphorylated amino acid and location in the protein, starting from the N-terminus. T – Threonine, S – Serine, Y – Tyrosine. The subunits of the PSI reaction centre psaA-psaE are colored bright green and labelled A-E respectively. The other subunits of the PSI reaction centre psaF—psaO are coloured dark green and labelled F-O. The LHCI subunits lhca1a, and lhca1-lhca9 are olive green coloured and numbered accordingly. See also Table 1 and Table S1. The location on the scheme is approximate.

**Figure 4.**
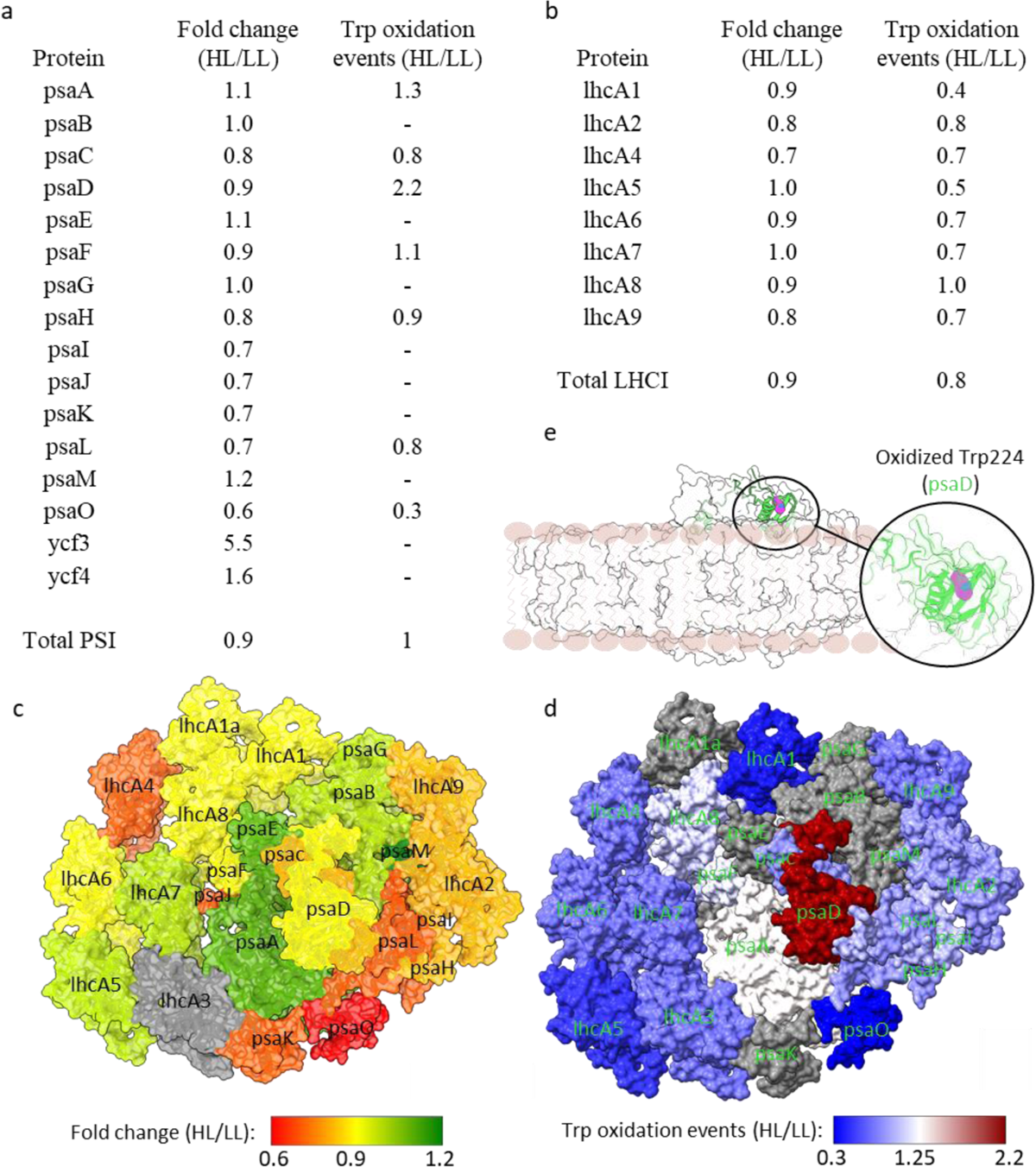
The number of oxidized tryptophans in psaO is reduced while those of psaD are increased in HL grown cells. (a + b) The fold change of PSI and LHCI subunits in LL and HL grown cells. Fold change was calculated by dividing the sum of peptides detected across three biological repeats of PSI-LHCI purified from HL grown cells by the sum of those detected in LL grown cells. See also Table S3. The same follows for the calculation of tryptophan oxidation events, in which peptides harbouring oxidized tryptophan were detected. see also Table S5. (c) Graphical representation of PSI-LHCI subunits fold-change values from panels a and b. Grey coloured represents the lncA3 subunit that was not detected in this analysis. (d) Graphical representation of fold-change values of oxidized tryptophan in PSI-LHCI subunits from panels a and b. Grey coloured subunits did not contain oxidized tryptophan. (e) Oxidative damaged tryptophan of psaD is shown in magenta, with PSI-LHCI complex as background (side view). For full details see Tables S3 and S4. Compelexes adapted from adapted from PDB: 6ZZX.

**Figure 5.**
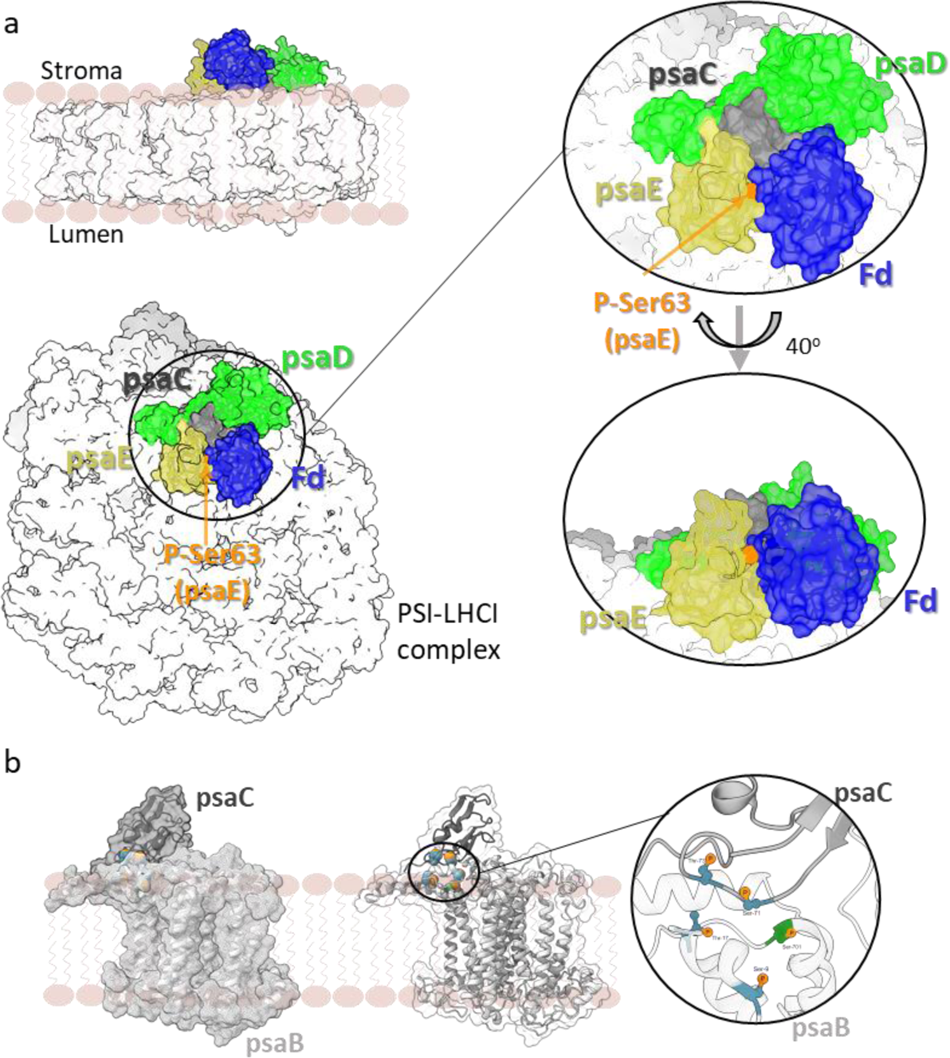
Phosphorylation at the psaB-psaC and psaE-Fd interfaces. (a) side and diagonal view of the PSI-LHCI complex. Phosphorylated Ser63 at the interface of psaE with Fd is indicated. psaC, psaD, and pseE form the binding site of Fd. (b) psaB and psaC are phosphorylated at their interface as indicated by orange orbs. Blue backbone indicates accumulation in LL conditions and green backbone indicates accumulation in HL conditions. Other PSI-LHCI subunits were removed for clarity.

### psaE and psaO are phosphorylated in LL-grown cells, lhcA6 is more phosphorylated in HL-grown cells

We found that some proteins were more phosphorylated under HL or LL growth conditions, and some proteins were phosphorylated to a similar extent under both conditions, but at different sites (Fig. 3, Table 1, Fig S1). psaE, psaO, psaC, psaF, lhcA7, and lhcA8 were phosphorylated in LL conditions. In psaB, 6 phosphorylated sites were detected, 3 of which were phosphorylated in LL- and 3 in HL-grown cells. psaK showed a similar pattern, with 2 sites phosphorylated in LL- and 4 in HL-grown cells. lhcA6 was found to be heavily phosphorylated under both conditions but to a larger extent in HL (Fig. 3 and Table 1). Phosphorylated psaA peptide was only detected in HL samples. Similarly, ycf4, an enzyme crucial for PSI assembly, was found phosphorylated exclusively under HL growth conditions. lhcA2 maintained the same levels of phosphorylation under both light conditions. These results suggest that light growth intensity can regulate the location and extent of phosphorylation of at least some of the PSI proteins.

### Phosphorylation of amino acids located in the psaB-psaC and psaE-Fd interfaces

Analysis of the localization of phosphorylated amino acids revealed that under LL growth conditions PSI-LHCI was phosphorylated exclusively at amino acids at the stromal side of the thylakoid membrane, while under HL growth conditions, the localization varied. psaB and psaK contained stromal-facing phosphorylation under LL, but luminal-facing (psaK), or phosphorylation in the hydrophobic transmembrane helix (psaB) in HL cells (Fig. 3). In both this and our previous work (Levin *et al*., 2023), we detected 5 phospho-sites in PSI subunits that are involved in PSI-Fd binding, specifically, 2 sites in psaC (Ser71 and Thr73) and 3 sites in psaE (Thr58, Ser86 and Ser63) (Figs. 2 and 5a and Table 1). Notably, the phosphorylation of Ser63 in psaE, identified in LL, but not HL *C. ohadii* cells in our earlier research, is located directly at the interface with Fd (Fig. 5a). The phosphate on Ser63 is expected to negatively charge the serine, possibly influencing psaE-Fd interactions. Interestingly, Ser63 is conserved in cyanobacteria and green algae, but not plants and red algae (Fig. S2). psaB and psaC are also phosphorylated at their interface, where a cluster of 5 phosphor-sites is located. These phosphorylations are expected to have a substantial impact on psaB and psaC binding to each other and perhaps on their stability (Fig. 5b). The 4 psaK phosphorylation sites exclusively observed in HL cells were closely located and formed a cluster of phosphorylated amino acids on the luminal side of the membrane (Fig. 3 and Table 1). lhca6 and lhca7, which were extensively phosphorylated under both HL and LL conditions, were phosphorylated only on the stromal side (Figs. 2a and 3).

### Phosphorylation of Ser63 in psaE may stabilize PSI binding to ferredoxin

To study the influence of Ser63 phosphorylation in PsaE on the stability of the PsaC-PsaD-PsaE complex with ferredoxin (Fig. 6a), we performed comparative molecular dynamics simulations and examined the structural characteristics of the systems. We found that in the unphosphorylated state, Ser63 primarily interacts with Val74 within the backbone (Fig. 6b), while its hydroxyl group in the side chain rarely forms interactions with Arg41 in PsaE and with amino acids in ferredoxin, such as Arg43, Glu32, and Tyr40, collectively accounting for less than 2% of the total simulation time (Table S4). Depending on the orientation of the phosphate group in Sep63, we identified two stable conformational states. In the first state (Fig. 6C), Sep63 interacts predominantly with Arg41 in the PsaE protein. Specific donor-acceptor interactions with Arg43 and Glu32 in ferredoxin are minimal, comprising less than 1% of the sampling. Nonetheless, there is a noticeable stabilization of the phosphorylated serine’s position relative to the side chain of Arg43 (Fig. 7), indicating a nonspecific electrostatic attraction between Arg43 and Sep63. In the second conformational state (Fig. 6D), Sep63 retains its ability to interact with Arg41 in the PsaE protein, but more importantly, it simultaneously establishes strong and specific donor-acceptor interactions with amino acids in ferredoxin, such as Arg43, Glu32, and Tyr40, with the primary interaction being with Arg43 (Table S4). It is plausible that the first conformational state represents the initial stage of complex formation with ferredoxin, while the second state reflects the result of its induced fit.

**Figure 6.**
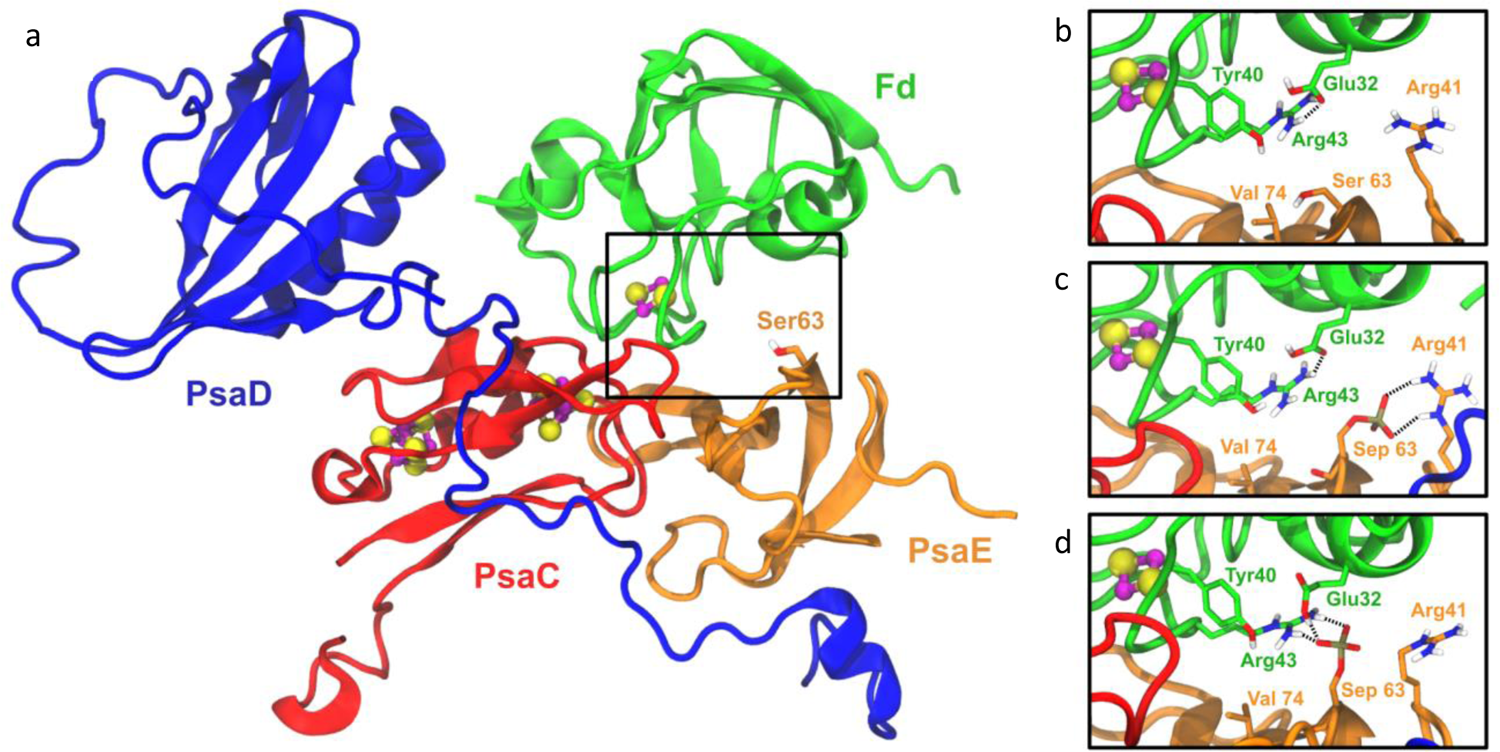
Ribbon model of the PsaC-PsaD-PsaE complex with ferredoxin (Fd) (a) and the interactions near Ser63 within the unphosphorylated (b) and phosphorylated conformations 1 and 2 (c and d) of the PsaE protein. The short-range donor-acceptor interactions are marked by black dashed lines.

**Figure 7.**
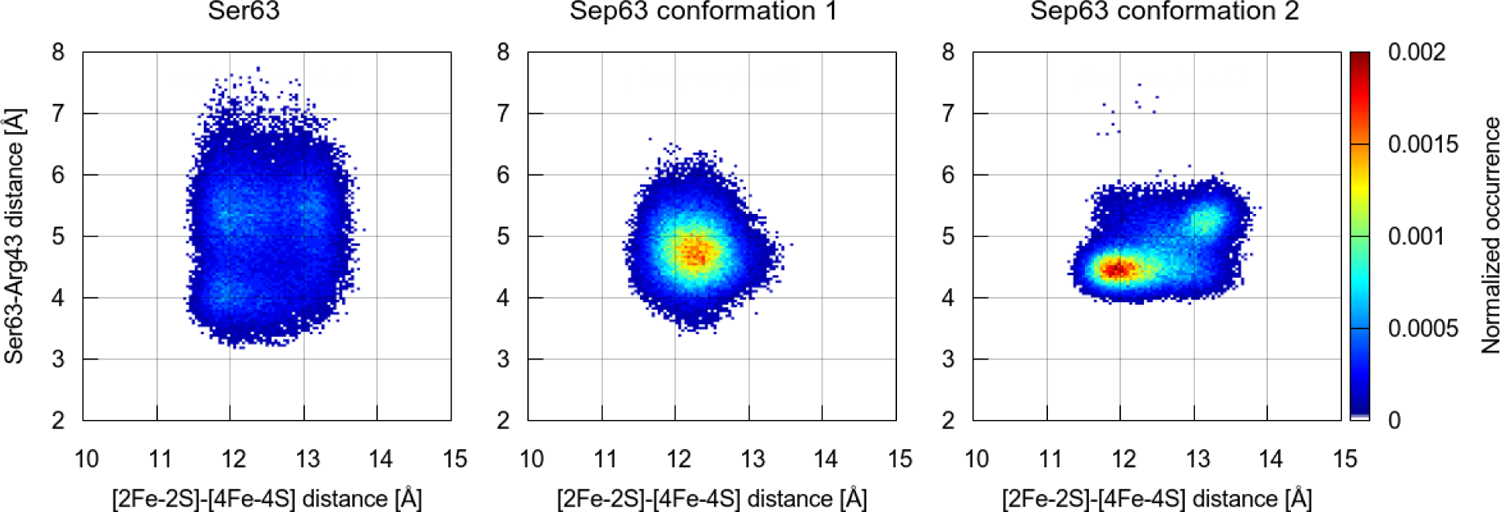
Relationship of the Ser63-Arg43 distance and [2Fe-2S]-[4Fe-4S] distance in the unphosphorylated (Ser) and phosphorylated (Sep) states of the PsaE protein.

### psaO levels are markedly reduced in PSI-LHCI of HL-grown cells

We assessed the relative abundance changes of the PSI-LHCI subunits in the unenriched samples to monitor possible adaptation to HL or photodamage. Among the 14 PSI and 8 LHCI subunits detected in the MS analysis, the levels of psaO were most significantly decreased in HL PSI-LHCI, presenting a fold-change value of 0.6 when compared to LL cells (Fig. 4 and Table S3). In a recent report detailing the structure of PSI-LHCI of LL-grown *C. ohadii*, psaO was found to reside on the periphery of the PSI complex but was absent in PSI-LHCI of HL *C. ohadii* cells (Caspy *et al*., 2021). psaO functions in binding LHCII subunits during state transitions in algae and plants (Jensen *et al*., 2004; Pan *et al*., 2018). However, HL grown *C. ohadii* cells were found to lack state transitions and undergo only negligible PSII protein phosphorylation, and therefore, binding of LHCII to PSI does not occur (Levin *et al*., 2023). This may be explained by the evident reduction of psaO levels in HL-grown cells. psaK and psaL, two additional subunits that are involved in state transitions, showed a slightly higher 0.7-fold change value compared to LL PSI, as did psaI and psaJ. The subunits that reside in close proximity to the photosynthetic core maintained a similar abundance in LL vs. HL conditions, as indicated by a fold-change of close to 1.0. psaA, psaB, psaC, psaD, psaE, psaF, and psaM exhibited fold-change values of 0.8-1.2 in HL cells, as did the peripheral units psaG and psaH (Fig. 4a and 4c and table S3). Similarly, all lhcA1-9 subunits, excluding lhcA3, which was not detected in the analysis, showed fold-change values of 0.7-1 (Fig. 4b and 4c and Table S3).

### psaD accumulates oxidized tryptophan while in LHCI the oxidation levels are reduced in HL PS-LHCI

Although PSI is far less sensitive to photodamage when compared to PSII, under certain conditions, it may undergo photoinhibition due to the accumulation of reactive oxygen species (ROS) (Sonoike, 2011; Lima-Melo *et al*., 2021). To determine whether HL induces oxidative stress to the PSI-LHCI subunits of *C. ohadii*, an analysis of oxidized tryptophan residues was conducted (Fig 1e). Intriguingly, oxidized tryptophans were detected in all LHCI subunits. However, all except one accumulated reduced levels of oxidized tryptophan in HL compared to LL cells. lhcA1 and lhcA5 accumulated 0.4 and 0.5 of the amounts of oxidized tryptophans compared to LL cells. For lhcA4, lhcA6, lhcA7, and lhcA9 it was 0.7, lhcA2 was 0.8, and lhcA8 was 1. No LHCI subunit accumulated more oxidized tryptophans in HL compared to LL cells (Fig. 4b and 4d and Table S5). Since the trytophans are oxidized by ROSs, the less accumulation in HL cells suggests the utilization of an efficient quenching mechanisms that reduce ROS accumulation in the LHCI of HL-grown cells. These quenching mechanisms are probably developed during the adaptation of the LL to HL cells (Levin *et al*., 2023).

In PSI, oxidized tryptophans were detected in 7 subunits (Fig. 4a and 4d and Table S5). Unlike the LHCI subunits, the PSI subunits seem to generally maintain similar levels of oxidized tryptophans in LL and Hl cells. psaC, psaF, psaH, and psaL show fold change values ranging between 0.8-1.1 in HL compared to LL cells. Oxidized tryptophan in psaA is slightly more accumulated in HL cells with a fold change value of 1.3. Interestingly, psaD, which was oxidized on a single tryptophan at position 224 (Fig. 4e), showed a 2.2-fold increase in its oxidation in HL cells, suggesting more sensitivity to oxidative stress of this subunit in HL cells (Fig. 4a and 4d and table S5). On the contrary, psaO tryptophan oxidation was markedly reduced in HL cells with a 0.3-fold change compared to LL cells. However, it is important to note that psaO levels are significantly lower in HL PSI, resulting in fewer psaO peptides that are available for oxidation. Further, we investigated whether the oxidation of psaD results in protein degradation. Semi-specific peptide searches facilitate the detection of endogenous proteolytic activity, often evident in protein degradation processes. Such activity leads to protein cleavage, producing truncated versions with terminal amino acids that may differ from the specificity of the enzyme used during proteomic sample processing (Fahrner *et al*., 2021). In our search for semi-tryptic-cleaved peptides in the PSI of both LL and HL cells as potential indicators of protein degradation, we observed no increase in peptide cleavage for psaD or any other PSI subunit at HL cells (Table S6).

## Discussion

This work showed that *Chlorella ohadii,* despite its limited state transitions and PSII protein phosphorylation, exhibits extensive phosphorylation in its LHCI proteins under both LL and HL growth conditions. Among all other PSI-LHCI subunits, the phosphorylated peptides of lhca6 and lhca7 were markedly overrepresented (Table 1). To date, there is no evidence suggesting that similar to LHCII, LHCI subunits detach from PSI or migrate to PSII in state I conditions. Therefore, it is unlikely that this phosphorylation initiates a state-transition-like process of the LHCI. In addition, no correlation was found between LHCI phosphorylation and oxidative damage (Table 1 and Fig. 4). However, the significant phosphorylation of lhca6 and lhca7 observed under growth conditions at different light intensities indicates its significant role in modulating light-harvesting activity under different light conditions. This modulation may be achieved by inducing structural changes and/or alteration in interactions with other antenna proteins tailored to the specific light-harvesting needs at a given light intensity. This hypothesis is supported by the observation that the phosphorylated amino acids are located at the interface with other LHCI subunits (Fig. 2).

Protein phosphorylation of PSII has been extensively studied and its protection roles in response to photodamage were extensively studied (Järvi *et al*., 2015; Theis & Schroda, 2016). In contrast, PSI is by far less sensitive to light-induced photoinhibition. Thus, it is likely that the role of PSI phosphorylation does not serve a similar purpose. In line with this, only the psaD subunit exhibited accumulated oxidative damage in tryptophan under HL conditions (Fig 4a). This was despite the absence of psaD phosphorylation or reduction in quantity (Table 1).

Alternatively, PSI subunit phosphorylation may transiently occur during the PSI complex biogenesis and/or disassembly and degradation processes. It should be emphasized that the current proteomics analysis was performed on purified PSI complexes that were isolated from thylakoids. Therefore, if the phosphorylation occurs during the assembly or disassembly of PSI while it is not active in the electron transport, it will still be present in the whole PSI complex. Nevertheless, phosphorylation seems to occur in many subunits which play important functions in light harvesting and photosynthetic electron transfer in the PSI. Over the years, phosphorylations were detected in subunits involved in plastocyanin binding (psaF and psaN), Fd binding (psaC, psaD, and psaE), and state transitions (psaH, psaK, psaL, psaG) (Grieco *et al*., 2016). To the best of our knowledge, this is the first report of phosphorylation of psaO, which also has a role in state transitions.

Generating a PSI-LHCI phosphor-site map has enabled us to reveal that many of the phosphorylations occur at amino acids that are located at the interface with other PSI subunits, such as in psaB-psaC and psaE-Fd, suggesting a role in the regulation of their binding. We observed that the stabilization of interactions between phosphorylated Ser63 in psaE and Arg43 in Fd correlates with a slight reduction in the distance between the [2Fe-2S] cluster in Fd and the [4Fe-4S] cluster in the PsaC protein (Figure 7). Therefore, one could speculate that Ser63 phosphorylation may facilitate the electron transport from PsaC to ferredoxin by stabilizing the interaction between phosphorylated Ser63 in PsaE and Arg43 in ferredoxin, which directly neighbors Cys42 coordinating the [2Fe-2S] cluster. Additionally, stronger interactions between ferredoxin and PsaE may result in a prolonged association of ferredoxin and, consequently, may allow for an extended period for electron transport. However, it’s important to note that this hypothesis requires further validation due to the limitations of our model, including the absence of a complete PSI complex in its membranal environment and the classical treatment of iron-sulfur clusters.

Additionally, quantitative analyses of the PSI subunits found psaO levels to be most reduced in HL as compared to LL grown cells (Fig. 4a). Although no evidence for a relation between psaO phosphorylation in LL to its binding to the PSI complex is known, it is possible that under HL conditions, psaO is dephosphorylated before its detachment from PSI, as it was absent in the PSI-LHCI complex of HL grown *C. ohadii* (Caspy *et al*., 2021). The absence of psaO strongly reduces the capacity to perform state transitions (Zhang & Scheller, 2004; Jensen *et al*., 2004). The lack of state transitions in HL *C. ohadii* may be at least partially attributable to the reduction of psaO content under HL conditions. This mechanism may protect PSI from excess light, as it prevents the binding of additional LHC subunits. As no increase in oxidative damage was detected in psaO of HL cells (Fig. 4a), its reduced levels may indeed reflect an adaptive rather than a photodamaging process.

The exact mechanism of ROS-induced PSI photoinhibition is yet not fully understood (Lima-Melo *et al*., 2021). Although our analysis shows that most PSI-LHCI subunits of HL cells do not accumulate more tryptophans that undergo oxidative damage than LL PSI-LHCI, highlighting its strong resistance to photodamage, psaD seems to be the exception. In HL PSI-LHCI, psaD accumulates higher levels of oxidative damage to tryptophan, raising the possibility that psaD is a potential target for photodamage during HL-induced photoinhibition of PSI. Alternatively, psaD may act as a shield for the adjacent FeS clusters in psaC, which are thought to be the site of PSI photoinhibition (Lima-Melo *et al*., 2021), by having a higher affinity to ROS. Interestingly, despite growing in HL conditions which usually induce ROS production, the LHCI subunits of PSI-LHCI from HL-grown cells showed lower accumulation of oxidative damaged tryptophans. This suggests that adaptive mechanisms that are induced during the acclimation process of *C. ohadii* to HL are capable of protecting the LHCI subunits from oxidative damage.

In summary, this work demonstrated how growth light intensity affects the pattern and location of PSI-LHCI phosphorylation in *C. ohadii*. Notably, LHCI subunits (lhca6 and lhca7) undergo extensive phosphorylation which might be related to their light harvesting functions and interactions with LHCI subunits. While many other PSI-LHCI subunits are also phosphorylated, their degree of phosphorylation is markedly reduced, suggesting that these events are more frequent during the assembly/disassembly phases of the complex, rather than in the mature and active complex. The localization of the phosphorylation was often found on the interface of proteins, such as the Ser63 in the psaE interface with Fd, highlighting the possibility of its role in complex binding and stabilization and/or in energy transfer (Nishi *et al*., 2011; Younas *et al*., 2023). Moreover, we show that the LHCI of HL-grown cells is better protected from ROS and that psaD suffers oxidative damage under HL conditions.

## Acknowledgments

We thank the Smoler Proteomics Center at the Technion for performing the mass spectrometry experiments. We enormously thank Dr. Oded Kleifeld and Rawad Hannah for their fabulous help with analyzing the mass spectrometry data. Funding was provided by a “Nevet” grant from the Grand Technion Energy Program (GTEP), Israel Science Foundation grant (2199/22), and a Technion VPR Berman Grant for Energy Research.

## Author contributions

GL and GS designed the research and wrote the article. GL, MY, and GS performed the research and experiments. TP and FG performed the molecular dynamics analysis. NY assisted in generating the figures and provided consultation for data interpretation.

## Data availability

All the analyzed data that is discussed is included in this paper and its supplemented data. Unedited excel files that contain all of the detected peptides in all of the proteomics experiments that are discussed in this paper are provided as Supporting Datasets 1-5. RAW files for proteomics experiments are available upon request from the corresponding author at

## Supporting information

**Fig. S1** Isolation of PSI-LHCI complexes.

**Fig. S2** The phosphorylation site at Ser63 of psaE is not conserved in red alga and plants.

**Table S1** PSI-LHCI proteins are differentially phosphorylated in LL and HL-grown *C. ohadii* cells.

**Table S2** Total phosphorylated PSI-LHCI peptides that were detected (Provided as separate Excel file)

**Table S3** Total PSI proteins that were detected (Provided as separate Excel file)

**Table S4** Interactions of phosphorylated Ser63 of psaE with ferredoxin

**Table S5** Detected oxidized tryptophan (Provided as a separate Excel file)

**Table S6** Detected cleaved PSI-LHCI peptides (excluding trypsin cleaved peptides) (Provided as a separate Excel file)

**Dataset S1** Total proteins in untreated samples (Excel file)

**Dataset S2** Total peptides in untreated samples (Excel file)

**Dataset S3** Phosphorylated peptides in untreated samples (Excel file)

**Dataset S4** Phosphorylated peptides in Phospho-enriched samples (Excel file)

**Dataset S5** Oxidized peptides in untreated samples (Excel file)

