## Supporting information for "The protein phosphorylation landscape in photosystem I of the desert algae *Chlorella sp*"

The following Supporting Information is available for this article:

**Fig. S1** Isolation of PSI-LHCI complexes.

**Fig. S2** The phosphorylation site at Ser63 of psaE is not conserved in red alga and plants.

**Table S1** PSI-LHCI proteins are differentially phosphorylated in LL and HL-grown *C. ohadii* cells.

**Table S2** Total phosphorylated PSI-LHCI peptides (Provided as separate Excel file)

**Table S3** Total PSI proteins (Provided as separate Excel file)

**Table S4** Interactions of phosphorylated Ser63 with ferredoxin

**Table S5** detected oxidized tryptophan (Provided as a separate Excel file)

**Table S6** detected cleaved PSI-LHCI peptides (excluding trypsin cleaved peptides) (Provided as a separate Excel file)

**Dataset S1** Total proteins in untreated samples (Excel file)

**Dataset S2** Total peptides in untreated samples (Excel file)

**Dataset S3** Phosphorylated peptides in untreated samples (Excel file)

**Dataset S4** Phosphorylated peptides in Phospho-enriched samples (Excel file)

**Dataset S5** Oxidized peptides in untreated samples (Excel file)

**Fig. S1 Isolation of PSI-LHCI complexes.** Thylakoid membranes were isolated from low or high-light (LL and HL) adapted cells and were solubilized with α-DDM. Sucrose density gradient (SDG) was used to separate native thylakoid protein complexes. The PSI-LHCI complex (indicated) containing fraction was collected and underwent mass spectrometry (MS) analysis for the detection of total and phosphorylated peptides of the PSI-LHCI proteins (See figure 1 and materials and methods).

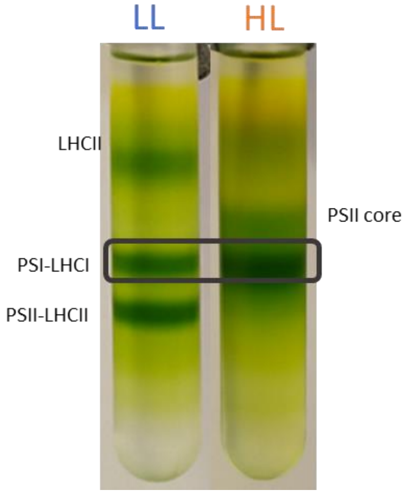


**Fig. S2 The phosphorylated Serine (63) of *Chlorella ohadii* psaE is conserved in green alga and cyanobacteria but not in red algae and plants.** Protein sequence alignment was performed using t-coffee multiple sequence alignment. The sequences of the proteins were obtained from the UniProt database. Conserved serine indicated by a green square. Arginine that were reported to interact with Fd are indicated by black squares.

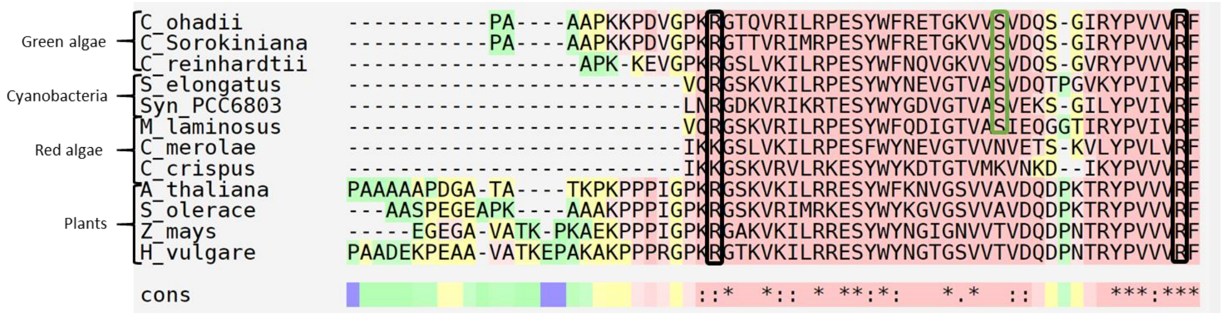

**Table S1** PSI-LHCI proteins are differentially phosphorylated in LL and HL grown C. ohadii cells. The phosphorylated residues are indicated by lower case blue coloured letters, as assigned by MSfragger software. Alternate locations for phosphorylation are indicated by lower case black letters. Peptide position and confidence of the position are indicated. Values in the 'P. peptide' columns represent the number of the phosphorylated peptides quantity as detected across 3 biological repeats for each light condition. The colour of the values indicates the sample in which the phosphorylated peptide was detected. Blue: phospho-enriched sample. Black: non phospho-enriched sample. Values in the 'Non-P. peptide' columns represent the number of the non-phosphorylated peptides quantity as detected across 3 biological repeats for each light condition in the non phospho-enriched sample. For full details see Table S2.

| Uniprot ID | Gene | Modified sequence | Residue # | P. location confidence (%) | P. peptides HL | P. peptides LL | Non-P. peptides: HL | Non-P. peptides: LL |
|---|---|---|---|---|---|---|---|---|
| P56341 | psaA | MtISPPER | T2 | 94 | 1 | 0 | 0 | 0 |
| W8SUA3 | psaB | EILEAQTPPsGsLGAGHK | S312 | 49 (49) | 0 | 1 | 26 | 30 |
| | | FSQALAQDPttR | T17 | 49 (49) | 0 | 1 | 19 | 33 |
| | | FSQALAQDPttRR | T17 | 49 (49) | 0 | 1 | 6 | 6 |
| | | FsQALAQDPTTR | S9 | 95-96 | 0 | 2 | 19 | 33 |
| | | PVALsIVQARLVGLtHFsVGYVLTYAAFLIASTSGKFG | S701, T711, S714 | 90, 90, 90 | 1 | 0 | 3 | 4 |
| A0A2I4S6V8 | psaC | VYLGSEtTR | T73 | 83-95 | 1 | 3 | 16 | 15 |
| | | VYLGsETTR | S71 | 90-94 | 4 | 12 | 16 | 15 |
| A0A2P6U4S6 | psaE | EtGKVVSVDQSGIR | T58 | 97 | 0 | 1 | 0 | 1 |
| | | VNYAGVStNNyALDEVEEVKA | S86 | 47 (47) | 0 | 1 | 118 | 111 |
| | | ETGKVVsVDQSGIRYPVVVR | S63 | 97 | 0 | 1 | - | |
| A0A2P6TPV8 | psaF | NGsLMEDDAK | S469 | 100 | 0 | 5 | 50 | 59 |
| | | NGsLMEDDAK | S469 | 100 | 2 | 8 | 50 | 59 |
| A0A2P6U0J1 | psaK | GAVVVRADYLGsttNQIMVLSTFLPLVAGR | S35, T36 | 63, 63 (63) | 1 | 0 | 2 | 0 |
| | | GAVVVRADyLGsttNQIMVLSTFLPLVAGR | Y32 | 46, 59 (46) (46) | 1 | 0 | 2 | 0 |
| | | FGLAPTstR | T59 | 45 (45) | 0 | 1 | 8 | 13 |
| | | FGLAPtstR | S60 | 33 (33) (33) | 0 | 1 | 8 | 13 |
| A0A2P6THB2 | psaO | GPsAWGAyEEPLsLVAGFLGWFAPSNIK | S45 | 33 (33) (33) | 0 | 1 | 1 | 5 |
| E1ZHX0 | lhcA2 | HLDGsMLGDYGFDPLR | S48 | 97 | 1 | 1 | 1 | 1 |
| A0A2P6TPR7 | lhcA6 | KPGsMDQDPIFSNFK | S145 | 95 | 49 | 29 | 46 | 64 |
| | lhcA7 | NPGSQGsGILPEDFK | S127 | 89 | 0 | 6 | 24 | 29 |
| | | NPGsQGSGILPEDFK | S124 | 94 | 4 | 18 | 24 | 29 |
| | | tVGGPYVGGR | T144 | 97 | 5 | 14 | 13 | 16 |
| A0A2P6TZ50 | lhcA8 | GTGIsGyPGGAFNPFGLGNSSK | S167 | 49 (49) | 0 | 1 | 0 | 1 |
| W8SUD8 | ycf4 | MtAQEAIIR | T2 | 100 | 2 | 0 | 0 | 0 |

**Table S2** Total detected phosphorylated peptides of PSI-LHCI are reported for both Phospho-enriched and non-phospho-enriched samples. Similar to Table S1, but with additional details. Provided as a separate Excel file.

**Table S3** Total PSI proteins that were detected and identified using non-phospho-enriched samples. Provided as a separate Excel file.

**Table S4** Normalized occurrence of the short-range donor-acceptor interactions of Ser63 in the unphosphorylated and phosphorylated states of the PsaE protein. Specific interactions are listed in the following format: acceptor residue-acceptor atom--donor residue-donor atom.

| **Ser63 state / Interactions** | **Ser63** | | **Sep63 conformation 1** | | **Sep63 conformation 2** | |
|---|---|---|---|---|---|---|
| **Ser63 – Val74**<br>(PsaE-PsaE) | S-O--V-N<br>V-O--S-N | **0.6549**<br>0.0153 | - | | S-O--V-N | 0.2037 |
| **Ser63 – Arg41**<br>(PsaE-PsaE) | S-OG--R-NE<br>S-OG--R-NH2<br>S-OG--R-NH1<br>S-OG--R-NH1<br>R-NH2--S-OG | 0.0010<br>0.0073<br>0.0001<br>0.0001<br>0.0001 | S-O1P--R-NH2<br>S-O2P--R-NE<br>S-O2P--R-NH2<br>S-O1P--R-NE<br>S-O3P--R-NE<br>S-O3P--R-NH2<br>S-OG--R-NH2 | **0.6900**<br>**0.6244**<br>0.0795<br>0.0111<br>0.0033<br>0.0029<br>0.0001 | S-O1P--R-<br>NH2 S-O2P--<br>R-NH2<br>S-O1P--R-NE<br>S-O2P--R-NE<br>S-O1P--R-<br>NH1<br>S-O2P--R-<br>NH1<br>S-OG--R-NH1<br>S-OG--R-NE<br>S-OG--R-NH2 | 0.3570<br>0.2510<br>0.1790<br>0.1321<br>0.0430<br>0.0181<br>0.0015<br>0.0271<br>0.0007 |
| **Ser63 – Arg43**<br>(PsaE-Fd) | S-OG--R-NH1<br>R-NH1--S-OG<br>S-OG--R-NH1<br>S-OG--R-NE | 0.0038<br>0.0001<br>0.0001<br>0.0002 | S-O1P--R-NH1<br>S-O3P--R-NH1<br>S-O1P--R-NH2<br>S-OG--R-NH1<br>S-O3P--R-NH2 | 0.0006<br>0.0005<br>0.0004<br>0.0003<br>0.0001 | S-O3P--R-<br>NH1<br>S-O3P--R-<br>NH2<br>S-O2P--R-<br>NH2<br>S-O2P--R-<br>NH1 | **0.7337**<br>0.0926<br>0.3141<br>0.0170 |
| **Ser63 – Glu32**<br>(PsaE-Fd) | E-OE2--S-OG<br>E-OE1--S-OG | 0.0019<br>0.0001 | S-O1P--E-OE2 | 0.0050 | S-O3P--E-OE2<br>S-O2P--E-OE2 | **0.6958**<br>0.1399 |
| **Ser63 – Tyr40**<br>(PsaE-Fd) | Y-OH--S-OG<br>S-OG--Y-OH | 0.0004<br>0.0001 | - | | S-O3P--Y-OH | 0.1529 |

**Table S5** detected oxidized tryptophan in PSI-LHCI from non-phospho-enriched samples.

Provided as a separate Excel file.

**Table S6** detected cleaved PSI-LHCI peptides in non-phospho-enriched samples. Only peptides that were cleaved by proteases other than trypsin were taken into consideration. Peptide cleavage may indicate protein degradation. Provided as a separate Excel file.

**Dataset S1** Total proteins in untreated samples (Excel file)

**Dataset S2** Total peptides in untreated samples (Excel file)

**Dataset S3** Phosphorylated peptides in untreated samples (Excel file)

**Dataset S4** Phosphorylated peptides in Phospho-enriched samples (Excel file)

**Dataset S5** Oxidized peptides in untreated samples (Excel file)
